# First Indications for Long-Term Benzodiazepine and Z-drugs use in the United Kingdom

**DOI:** 10.1101/085183

**Authors:** J. Davies, T. C. Rae, L. Montagu

## Abstract

Benzodiazepines and Z-drugs (BZDs), hypnotic drugs used for insomnia and anxiety, are prescribed millions of times a year in the UK. Although guidance from the relevant regulatory authorities (NICE and BNF) indicates them only for short-term use, the evidence suggests that many patients have been taking these drugs for much longer, often for decades. At present, there are no up-to-date, evidence-based estimates of the scale of long-term BZD use in the UK, which has prevented making a strong case for the need for withdrawal services. However, data obtained recently on BZD use from a number of GP surgeries (covering nearly 100,000 registered patients) in the North of England, allow such projections to be calculated. Scaling the results to a national level suggests that there are over a quarter of a million patients in the UK using BZDs for periods far longer than recommended. The projections also suggest that nearly half this number may be willing to accept help to stop their dependency on BZDs. These results indicate a serious problem, which should be addressed by more research into the harms associated with long-term BZD use, the provision of withdrawal services, and a national helpline to support patients with BZD dependency.

## Introduction

Benzodiazepines are hypnotic drugs that enhance the activity of gamma-aminobutyric acid (GABA) at the GABAA receptor. Zolpidem, zopiclone and zaleplon, commonly known as Z-drugs, are non-benzodiazepine hypnotics that share a similar mode of action but are chemically distinct.^1^ Both benzodiazepines and Z-drugs (BZDs) are indicated for the short-term relief of severe or disabling anxiety, whether this occurs alone or in association with insomnia or short-term psychosomatic, organic, or psychotic illness.^2^ Approximately 16 million prescriptions for BZDs were issued in England during 2015 – a figure that has broadly remained steady since 2011.^3^ Concerns regarding the addictive potential of these drugs have been highlighted for many years,^4 5^ leading the BNF to recommend that uninterrupted usage not exceed four weeks,^6^ as long-term use can cause adverse neurological, cognitive and physical effects, but also high degrees of physical and psychological dependency.^7 8 9^

It is now recognised that withdrawal from benzodiazepines and Z-drugs can be very protracted, generally lasting between 6 to 18 months after the last dose, and sometimes even longer.^10 11^ Withdrawal charities report numerous cases of patients taking at least three or four years to recover, and some are left with residual symptoms such as tinnitus, which can persist for years beyond this timeframe.^12^ In all, long-term BZD use (and withdrawal from it) can generate a range of long-term disabling effects, which can impact negatively on many aspects of a person’s life, threatening relationships, careers and financial stability.^13^

Despite previous claims by the Department of Health’s National Treatment Agency,^14^ there are currently no appropriate NHS services available to support these long-term BZD users during withdrawal, and instead patients have to rely on a small number of inadequately resourced specialist support charities, whose provision extends to only a handful of local regions. Consequently, in recent years a nationwide patient movement has materialised, alongside two separate All-Party Parliamentary Groups, which together have lobbied the Department of Health and Public Health England for the funding of specialist withdrawal services for those affected by prescribed drug dependence. A general response by each Department, however, has focused on the absence of authoritative data on the number of long-term users in the UK, and accordingly, on the number of those affected by dependence and withdrawal. In the absence of a robust indication of need, both departments have claimed it has been difficult to establish a clear basis for the provision of prescribed drug withdrawal services. This response betrays a common problem encountered by campaigners, who report unwillingness on the part of government departments and some senior members of the medical profession to accept the scale of the problem today.^15^ An example of this can be heard on the BBC Radio 4 ‘Face the Facts’ programme from 31 July 2011, where ex-president of the Royal College of General Practitioners, Dr Clare Gerada, describes the current situation with benzodiazepines as a ‘prescribing success story’ despite being given figures on the programme that demonstrate the opposite.^16^ For example, the Bridge Project withdrawal charity has produced data showing that the majority of users it has identified have been taking benzodiazepines for many years; and of the long-term users taking up their support services, over 76% had been taking these drugs for more than six years.

If withdrawal services are to be considered for government funding, an up-to-date, evidence-based estimate of the number of long-term BZD users is urgently required to inform policy decision-making and to justify the allocation of resources. The best existing estimate, extrapolated from a small survey by the BBC Panorama programme in 2001,^17^ is now 15 years out of date and therefore immaterial to current policy decision-making.^18^ Nonetheless, the continued existence of a large online community of prescribed drug dependents^19^ suggests that the numbers may be substantial. If viable services are therefore to be considered today, it is vital to determine the current levels of long-term BZD users in the general population. New data on the percentage of prescribed drug dependents from a sample of general practice surgeries allow such estimates to be calculated.

## Materials and Methods

The data are derived from a recent survey of GP surgeries in Bradford, UK, conducted by The Bridge Project. The information was accessed by submitting a request to each practice. The data include the number of registered patients at each surgery, the number of patients using BZDs, the number of patients whose use can be considered ‘long-term’ (defined as persons taking these medications for at least 12 months, which is significantly beyond the 2-4 weeks recommended by the BNF) and the number of long-term BZD users who agreed to accept help in ending their pharmacological dependency.

The data themselves are subject to several shortcomings, in terms of representativeness; in particular, they are taken only from a single geographical area, although spanning both urban and semi-rural areas. They have, however, been subject to a selection process, as the figures reported here exclude those under age 16 and over age 80, those in receipt of palliative care, those suffering illness at the time of the survey, those with a diagnosis of epilepsy, and those with severe and enduring mental health issues. Given these exclusion criteria, and the fact that the Bridge Project has encountered surgeries that appear to have underreported the number of BZD users, the projections reported here are necessarily conservative. Although still a relatively small sample, with all of the caveats that entails, these data represent the only reliable information available on long-term use of BZDs in the UK.

To estimate the total number of long-term BZD users in the UK, the mean percentage and standard error of long-term users across surgeries is calculated. These percentages values are converted into an estimate of national long-term BZD users through multiplying them by the number of patients aged 16-80 registered at UK GP surgeries (after applying the exclusion criteria listed above) from figures published by the Health and Social Care Information Centre (2014).^20^ An alternative estimate is calculated as the percentage of such users across the entire sample multiplied by the number of registered patients. The number of patients who might be willing to accept help to end long-term use is determined by multiplying the estimated number of long-term users by the overall percentage of BZD users who have agreed to take advantage of the charitable services offered at the surgeries sampled. All calculations are performed in MS Excel 2010.

## Results

After filtering, the surgeries surveyed have a total of 97,798 registered patients. The mean percentage of registered patients aged 16-80 classed as long-term (>1yr.) benzodiazepine and Z-drug users across the surgeries sampled is 0.69% ± 0.15%. This yields a mean projection of 296,929 ± 64,376 long-term users in the UK. Using the overall percentage of long-term users from all surgeries sampled yields a very similar estimate (266,905). In either case, the values indicate a substantial problem. In addition, we estimate that 119,165 of these patients (40.13% of the mean projection of UK long-term users) are likely to be willing to accept services designed to free them from prescribed drug dependency.

## Discussion

The results reported above indicate that a substantial number of people in the UK are likely to be taking highly dependency-forming benzodiazepine and Z-drug medication far beyond the recommended time scales. As there is already evidence that long-term use of BZDs causes adverse physiological and neurological affects,^21 22^ and protracted withdrawal,^23^ this represents a serious problem, particularly at a time when the NHS is experiencing large-scale austerity and efficiency restrictions.^24^

In June 2012, APPGITA (All-Party Parliamentary Group on Involuntary Tranquiliser Addiction) undertook a survey to determine the level of the provision of services for involuntary addiction to drugs like benzodiazepines and Z-drugs.^25^ This followed an assertion by the Department of Health’s National Treatment Agency that there were withdrawal services available in most of the country.^26^ However, this assertion was based on an NTA report that only surveyed existing illegal drug and alcohol treatment services, rather than services for those suffering from prescribed drug dependence. To determine levels of provision for the latter group, APPGITA contacted 149 Primary Care Trusts, asking what services were available in their area to support those suffering involuntary prescribed drug dependence. Of the 100 who responded, 83 primary care trusts acknowledged that they had no services, 11 had partial services while only 6 confirmed that they had services.^27^

In an effort to compensate for this dearth of NHS services, there are five charities providing local services for individuals withdrawing from prescription drugs - Addiction Dependency Solutions (Oldham); Battle Against Tranquilisers (Bristol); the Bridge Project (Bradford); Bristol and District Tranquiliser Project (Bristol); REST Minor Tranquiliser Service (London). Historically, these charities emerged from the efforts of single individuals, who, having recovered from dependency themselves, have lobbied local commissioning groups for services in their area. However, as these charities cover less than 5% of the country in terms of the population they serve, it seems a costly failing that there is no nationwide service provision, particularly given the large sums spent supporting illegal drug and alcohol addiction, and given the moral obligation to help individuals who have become dependent predominantly through NHS GP practices. Due to the increased delegation of funding to local authorities since 2013, these charities assert that what little state funding they have received is now also imperilled. This has led to increased levels of uncertainty regarding financing, and to at least one withdrawal charity (CITA in Liverpool) shutting its doors in 2014 due to changes in local funding priorities. In short, as it is unreasonable to assume or expect that more individuals will elect to lobby for services in other areas of the country, it is plausible to state that the only way for nationwide services to be guaranteed is if the government insists on a national response, ideally with support from the Department of Health.

Given the various challenges in establishing useful national provision, we make the following four recommendations, which we believe will help address the levels of long-term use in the UK, reducing the harms associated with dependence and withdrawal.

1. More research is required into the harms associated with long-term benzodiazepine and Z-drugs use, as well as the demographics and geography of long-term users.

While there is extensive testimony from individuals who have been harmed by these drugs, there has been very little systematic research in key areas. We need to determine what percentage of long-term BZD users is affected by negative and withdrawal effects, and how this correlates with dosage, length of use and withdrawal method. Furthermore, while this current study provides the first clinically-rooted estimate for the number of long-term BZD users in the UK, there are no data showing more detailed demographic and geographic usage trends. Such data will be crucial in guiding withdrawal outreach programmes, should current provision be up-scaled. With reports of symptoms such as tinnitus and nerve pain lasting many years, more research is also urgently needed into the physiological and neurological harms associated with long term use.

2. Reduce prescribing levels by ensuring adherence by doctors to existing guidelines for prescribing and withdrawal, and develop new guidelines where needed.

Many of the patients experiencing problems with prescribed medicines would have avoided the associated harms if their doctors had simply adhered to the prescribing guidelines that are already in place. As this study reveals, while BZDs are indicated for short-term use only in the BNF, a large cohort continue to take these drugs long-term, and withdrawal charities report many cases of new long-term prescriptions. Additionally, in the experience of the withdrawal charities, there appears to be a correlation between the severity of symptoms and the speed of withdrawal. The harm sometimes caused by cold turkey withdrawal or a rapid taper cannot be overstated; for many users this can lead to years of severe suffering accompanied by the loss of career, marriages and financial security.^28^ We need to ensure that doctors adhere to the tapering guidelines in the BNF for BZDs, and that they support the development of new guidelines for antidepressants, based on withdrawal charities’ experience and best practice.

3. Mandatory national provision of prescribed drug withdrawal services

Most patients are unaware of the risks of dependence and long-term effects, and therefore do not seek out services to help with withdrawal. GP practices must position to actively identify and contact long-term users. In Oldham, Bradford and Liverpool, withdrawal charities have had considerable success working with GP practices to identify, communicate with and ultimately help patients safely withdraw from their medications, and this model of provision should be extended across the country. As we have stated earlier, the provision of these services should be made mandatory to ensure that all patients across the UK are at the very least offered the support they need if they elect to withdraw.

4. Establishment of a national helpline

A national helpline and accompanying website for prescribed drug dependence, would provide an essential resource for patients, carers, families and doctors, delivering a low cost, yet effective national response to a recognised public health issue. A national helpline would also be the first step towards the provision of local specialist support services, as it would also enable the NHS to gather data on the scale of the problem and highlight gaps in current local service provision. A national helpline would also get around the unhelpful response of ministers and the Department of Health over recent years which have argued that – following the devolution of the NHS – this is now a matter for local authorities and Clinical Commissioning Groups.

## Footnotes

1 V. Kapil, J. L. Green, C. Le Lait, D. M. Wood, P. I. Dargan. Misuse of benzodiazepines and Z-drugs in the UK. *The British Journal of Psychiatry*, 205 (5) 407-408 (2014).

2 British National Formulary. 4.1 Hypnotics and Anxiolytics. (http://www.evidence.nhs.uk/formulary/bnf/current/4-central-nervous-system/41-hypnotics-and-anxiolytics), Accessed, Sep 2016.

3 HSCIS (2016) *Prescriptions Dispensed in the Community -2005-2015, Report*. Website: http://content.digital.nhs.uk/catalogue/PUB20664/pres-disp-com-eng-2005-15-rep.pdf. Accessed, Oct, 2016.

4 Committee on Safety of Medicines. Benzodiazepines, dependence and withdrawal symptoms. *Current Problems*, 21: 1–4 (1988).

5 Jones, I. R., Sullivan G. Physical dependence on Zopiclone: Case reports. *British Medical Journal*, 316:117 (1998).

6 British National Formulary. 4.1 *Hypnotics and Anxiolytics*. (http://www.evidence.nhs.uk/formulary/bnf/current/4-central-nervous-system/41-hypnotics-and-anxiolytics), Accessed Sep 2016.

7 Ayers, S., Baum, A., McManus, C., Newman, S., Wallston, K., Weinman, J., eds. *Cambridge Handbook of Psychology, Health and Medicine* (2nd ed.). Cambridge University Press. p. 677. (2007).

8 Ashton, C. H. *Benzodiazepines: How They Work and How to Withdraw*. Institute of Neuroscience, Newcastle University, Newcastle upon Tyne. (2002) See http://www.benzo.org.uk/manual/index.htm. Accessed, 22 February 2016.

9 Gudex, C. Adverse effects of Benzodiazepines, *Social Science and Medicines*, 33: 587-97 (1991).

10 British National Formulary. 4.1 *Hypnotics and Anxiolytics*. (http://www.evidence.nhs.uk/formulary/bnf/current/4-central-nervous-system/41-hypnotics-and-anxiolytics) (2015). Accessed Sep 2016.

11 Ashton, C. H. Benzodiazepine withdrawal: outcome in 50 patients. *Br J Addict*, 82: 665-71 (1987).

12 Ashton, C. H. *Benzodiazepines: How They Work and How to Withdraw*. Institute of Neuroscience, Newcastle University, Newcastle upon Tyne. (2002). See http://www.benzo.org.uk/manual/index.htm. Accessed 22 February 2016.

13 Council for Evidence-based Psychiatry. *BMA Call for Evidence: Involuntary Dependence to Prescription Medications: Response from the Council for Evidence-based Psychiatry (“CEP”).* Unpublished Report (2014).

14 NTA report: Addiction to Medicine, 2011, page 16, see http://www.nta.nhs.uk/uploads/addictiontomedicinesmay2011a.pdf

15 Council for Evidence-based Psychiatry. *BMA Call for Evidence: Involuntary Dependence to Prescription Medications: Response from the Council for Evidence-based Psychiatry (“CEP”).* Unpublished Report (2014).

16 Face the Facts. *Prescribed Addiction*. BBC Radio 4. (31^st^ July 2011). Website: http://www.bbc.co.uk/programmes/b012wxxw. Accessed Oct 2016.

17 BBC Panorama, The Tranquiliser Trap, 2001 (transcript: http://news.bbc.co.uk/hi/english/static/audio_video/programmes/panorama/transcripts/transcript_13_05_01.txt). Accessed, Aug 2016.

18 Since the early 2000s medical awareness has grown regarding the of dangers long-term benzodiazepine use, potentially reducing long-term use. Furthermore, as benzodiazepine prescribing was at its height in the 1960s, 1970s we would expect the number of long-term benzodiazepine users to have peaked around the 2000, reducing thereafter.

19 For example, see: www.benzo.org.uk, www.recovery-road.org and www.benzobuddies.org.

20 Health and Social Care Information Centre. *Number of patients registered at a GP practice* (2014). Website: https://data.gov.uk/dataset/numbers_of_patients_registered_at_a_gp_practice/resource/a4df09f4-3d03-49e8-8dfa-f36bc2d076b9 Accessed: June 2016.

21 Nielsen, M., Hansen, E.H. & Gøtzsche, P.C. What is the difference between dependence and withdrawal reactions? A comparison of benzodiazepines and selective serotonin re-uptake inhibitors. *Addiction*. 107(5):900-8. (2012) doi: 10.1111/j.1360-0443.2011.03686.

22 Ayers, S., Baum, A., McManus, C., Newman, S., Wallston, K., Weinman, J., (eds.). *Cambridge Handbook of Psychology, Health and Medicine* (2nd ed.). Cambridge University Press. (2007).

23 Ashton, C. H. Benzodiazepine withdrawal: outcome in 50 patients. *Br J Addict*, 82: 665-71 (1987).

24 Roberts, A, et al. A decade of austerity? The funding pressures facing the NHS from 2010/11 to 2021/22. Nuffield Trust (2012). Website: http://www.nuffieldtrust.org.uk/publications/decade-austerity-funding-pressures-facing-nhs. Accessed, Sep, 2016.

25 Perrot J. *Survey of PCTs Recording Provision of Services for Involuntary Tranquiliser Addiction for the All Party Parliamentary Group for Involuntary Tranquiliser Addiction*, (2012). Website: http://www.yumpu.com/en/document/view/11700252/survey-of-services-john-perrott-appgita. Accessed June 2016.

26 National Treatment Agency for Substance Misuse. Addiction to Medicine. Website: http://www.nta.nhs.uk/uploads/addictiontomedicinesmay2011a.pdf. p.16 (2011). Accessed June 2016.

27 Perrot J. *Survey of PCTs Recording Provision of Services for Involuntary Tranquiliser Addiction for the All Party Parliamentary Group for Involuntary Tranquiliser Addiction*, (2012). Website: http://www.yumpu.com/en/document/view/11700252/survey-of-services-john-perrott-appgita. Accessed June 2016.

28 Council for Evidence-based Psychiatry. *BMA Call for Evidence: Involuntary Dependence to Prescription Medications: Response from the Council for Evidence-based Psychiatry (“CEP”).* Unpublished Report (2014).

